# The evolution of extreme fertility defied ancestral gonadotropin mediated brain-reproduction tradeoff

**DOI:** 10.1101/2020.04.30.070078

**Authors:** Hagai Y Shpigler, Brian Herb, Jenny Drnevich, Mark Band, Gene E Robinson, Guy Bloch

## Abstract

Gonadotropic hormones coordinate processes in diverse tissues regulating animal reproductive physiology and behavior. Juvenile hormone (JH) is the ancient and most common gonadotropin in insects, but not in advanced eusocial honey bees and ants. To probe the evolutionary basis of this change, we combined endocrine manipulations, transcriptomics, and behavioral analyses to study JH regulated processes in a bumble bee showing an intermediate level of sociliality. We found that in the fat body, more JH-regulated genes were upregulated and enriched for metabolic and biosynthetic pathways. This transcriptomic pattern is consistent with earlier evidence that JH is the major gonadotropin in the bumble bee. In the brain, most JH-regulated genes were downregulated and enriched for protein turnover pathways. Brain ribosomal protein gene expression was similarly downregulated in dominant workers, which naturally have high JH titers. In other species, similar downregulation of protein turnover is found in aging brains or under stress, and is associated with compromised long-term memory and health. These findings suggest a previously unknown gonadotropin-mediated tradeoff. We did not find a similar downregulation of protein turnover pathways in the brain of honey bees in which JH is not a gonadotropin but rather regulates division of labor. These differences between JH effets in the bumble bee and in the advanced eusocial honey bee suggest that the evolution of advanced eusociality was associated with modifications in hormonal signaling supporting extended and extremely high fertility while reducing the ancient costs of high gonadotropin titers to the brain.

## Introduction

Animals tightly regulate reproduction to enhance their fitness. Hormones play key roles in this process by integrating relevant external and internal information and by coordinating biochemical and physiological processes across cells and tissues throughout the body (Christensen, et al. 2012). In insects, juvenile hormone (JH) is the most commonly known gonadotropin regulating vitellogenesis, oogenesis, and other processes that are associated with reproduction (Wyatt 1997). Given that JH functions as a gonadotropin in females of the more basal Hemimetabola and most Holometabola insects, it is commonly accepted that the regulation of reproduction is the ancient and conserved function of JH in adult insects (De Loof, et al. 2001; Riddiford 2012; Roy, et al. 2018).

However, it has long been puzzling that JH does not function as a gonadotropin in all insects (Cameron and Robinson 1990; West-Eberhard and Turillazzi 1996; Robinson and Vargo 1997). One of the most remarkable exceptions is in honey bees and several species of ants, in which JH does not regulate adult female fertility but is rather involved in the regulation of age-related division of labor among functionally sterile workers (Robinson and Vargo 1997; Hartfelder 2000; Bloch, et al. 2009). Addressing this question is crucial for understanding both the evolution of advanced eusociality and the evolution of gonadotropic hormones in general.

To study variability in socially-related JH functions, we focused on the best-studied bumble bee, *Bombus terrestris*. Bumble bees provide an excellent model system with which to address this issue because they show a simpler form of social organization relative to honey bees and ants (Wilson 1971), and JH is their major gonadotropic hormone (Röseler 1977; Röseler and Röseler 1978; Bloch, Borst, et al. 2000; Amsalem, et al. 2014; Shpigler, et al. 2014). JH coordinates processes in various tissues that are involved in *B. terrestris* reproduction including the fat body, ovaries, wax production, and exocrine gland activity (Shpigler, et al. 2010; Amsalem, et al. 2014; Shpigler, et al. 2014). JH apparently also regulates processes in the central nervous system because JH manipulations affect behaviors such as dominance and aggression (Amsalem, et al. 2014; Pandey, et al. 2019). The bumble bee *B. terrestris* thus provides an excellent contrast to honey bees to address the question of the evolutionary change in JH function associated with eusociality, but little is known on the molecular processes by which JH coordinates the various cells and tissues in this species.

Here we combine hormonal manipulations, behavioral observations, and RNA sequencing (RNAseq) of the brain and fat body tissues of *Bombus terrestris* workers. We further take advantage of two “natural experiments” in which we compared gene expression in bumble bees that naturally differ in JH titers. First, we analyzed the brain transcriptomes of orphan workers in which the dominant individual in a group typically has high JH titers compared with low-rank groupmates. Second, we compared our worker fat body transcriptomes to genes differentially expressed between queens and (pre-reproductive) gynes, which naturally have high and low JH titers, respectively. Our results revealed a previously unknown gonadotropin-mediated tradeoff in which the activation of reproductive pathways in various tissues is associated with a significant downregulation of brain pathways involved in protein synthesis and turnover. This finding suggests that high JH titers are costly to the brain because they reduce protein biosynthesis that is needed for processes such as long-term memory, synaptic plasticity, and brain maintenance. Consistent with our premise, we did not find a similar influence of JH on the brain in the honey bee in which JH is not a gonadotropin, but rather regulates the age-related division of labor among sterile workers.

## Results

### JH regulates the expression of numerous fat body and brain genes

Manipulating circulating JH levels by surgically removing the JH-producing corpora allata (CA) modified the expression of thousands of fat body and brain genes (Fig 1A and 2A, for fat body and brain respectively). In the fat body, 4408 transcripts were differentially expressed between the Allatectomy (CA-), Sham-operated (S), Control (C) and Replacement Therapy (CA- +JH) treatments (ANOVA, false discovery rate, FDR < 0.05, 8 bees per treatment). Hierarchical cluster analysis separated the pattern of gene expression transcriptome of bees with reduced JH titers (CA-) from that of the three other treatments (Fig. 1A, B). The replacement therapy treated bees differed from the CA- bees, but these two groups were yet distinct from the Sham and Control treatments. Complementary pairwise comparisons identified 4032, 3274 and 2005 genes that were differentially expressed (FDR < 0.05) between the CA- bees and those from the Control, Sham and the Replacement Therapy treatments, respectively (Fig 1B, left). A subset of 1512 DEGs differed between the CA- treatment and all three other treatments. This overlap was significantly higher than expected by chance (χ^2^ _(df =7)_ = 8316; p << 0.001). The direction of the change (i.e., up- or down-regulation) of these shared genes was significantly positively correlated across all transcripts for the three comparisons (Fig. 1C, Supp Table 1A). Of these, a subset of 873 out of 1512 (57%) DEGs were upregulated and 639 (43%) were downregulated. An additional 1553 DEGs differed between the CA- bees and only two of the three treatment groups and are also likely to be regulated by JH (Supp Table 2).

**Figure 1.**
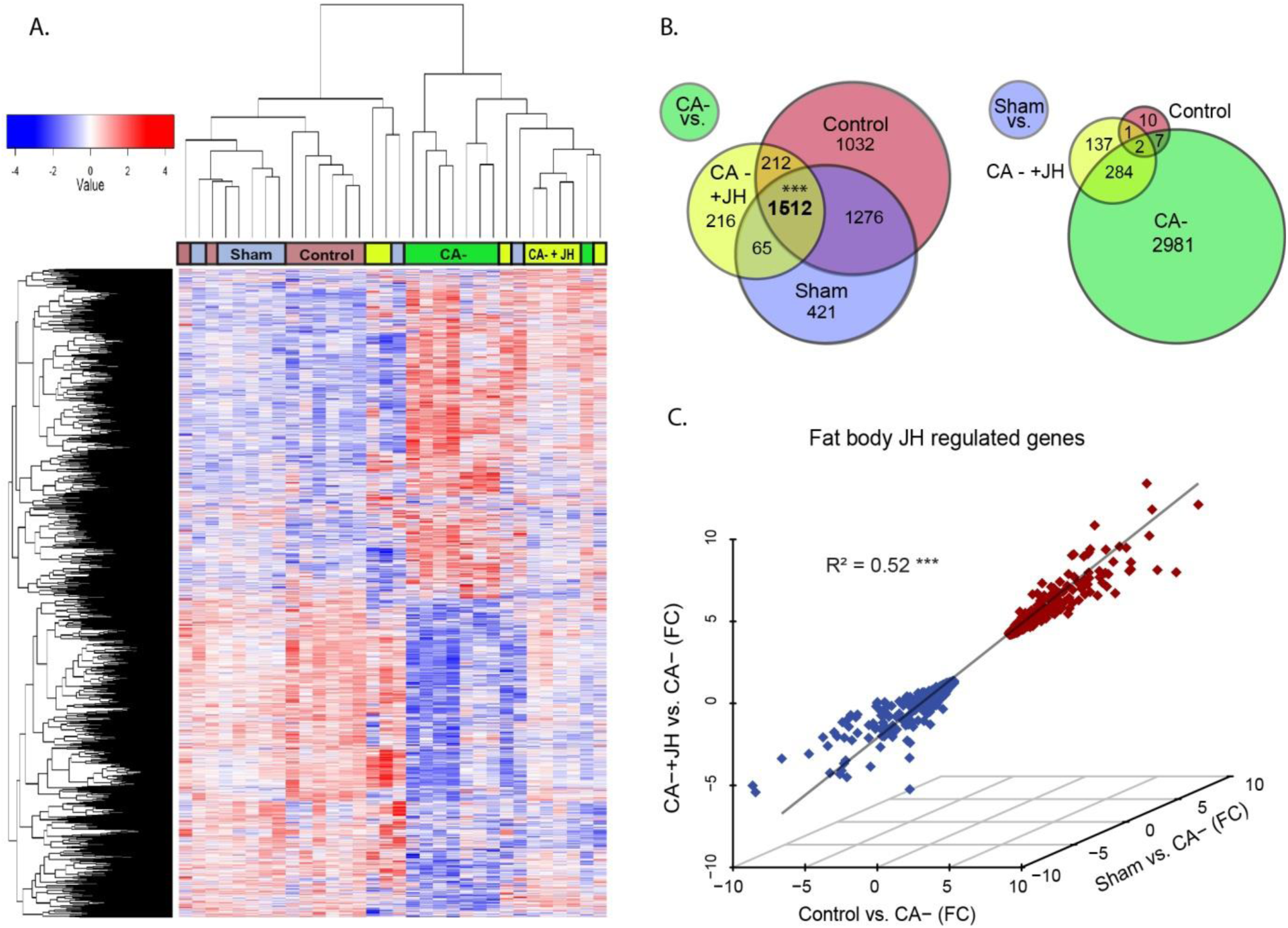
The effect of JH on gene expression in the fat body. Fat body transcriptomic analysis for bees subjected to a reduction in JH level (allatectomized, CA-, green), replacement therapy with natural JH (CA- + JH, yellow), handling control (*Control*, pink), and sham-operated bees (*Sham*, blue)(n=8 bees/treatment). **(A)** A heat map of 4032 DEGs (FDR p < 0.05) in the fat body. Each row represents one gene and each column an individual bee sample. The color scale represents expression level across all samples with greater expression represented in red and lesser expression by blue relative to the mean. **(B)** Venn diagrams comparing DEGs between the CA- treatment (left), or the Sham treatment (right) and the three other treatments. *** denotes a highly significant (p << 0.0001) overlap. **(C)** Three-dimension correlation analysis for the direction and fold expression for 1512 transcripts that were differentially expressed between the CA- and the three other treatments. Each diamond represents one DEG, red - upregulated by JH; blue - downregulated by JH. The R^2^ was obtained from a Person correlation test; *** - < 0.0001 (See Supp Table 1A for more details).

In the brain, JH manipulation affected the expression of 3060 transcripts. As in the fat body, hierarchical cluster analysis clearly separated the allatecomized (CA-) bees from bees subjected to the three other treatments (Fig. 2A). Importantly, the replacement therapy treatment was most similar to that of the sham-treated bees, which means that in also in the brain the operation itself had little or no effect on gene expression. Complementary pairwise comparisons revealed 2615, 2100 and 1195 transcripts differentially expressed (FDR p < 0.05), between the CA- bees and the Control, Sham, or Replacement Therapy treated bees, respectively (Fig 2B left). Seven hundred DEGs differed between the CA- bees and all three other treatments, which is significantly higher overlap than expected by chance (χ^2^ _(df =7)_ = 7425; p << 0.001, Fig 2B left). These 700 hundred genes provide a conserved estimation of JH regulated genes, of which 280 (40%) were up- and 420 (60%) down-regulated by JH. This pattern is significantly different from the fat body in which most genes are upregulated by JH (Z-test for proportions, Z = 7.44; p = 1.66E- 13). The direction of expression of the shared genes was significantly positively correlated and consistent across all transcripts for the three comparisons (Fig. 2C, Supp table 1B). An additional 1096 DEGs differed between the CA- bees and only two of the three treatment groups and are also likely to be regulated by JH (Supp Table 3). The strong effect of JH removal and the reverted pattern in bees subjected to replacement therapy show that JH regulates the expression of many genes in both the brain and the fat body of bumble bee workers.

**Figure 2.**
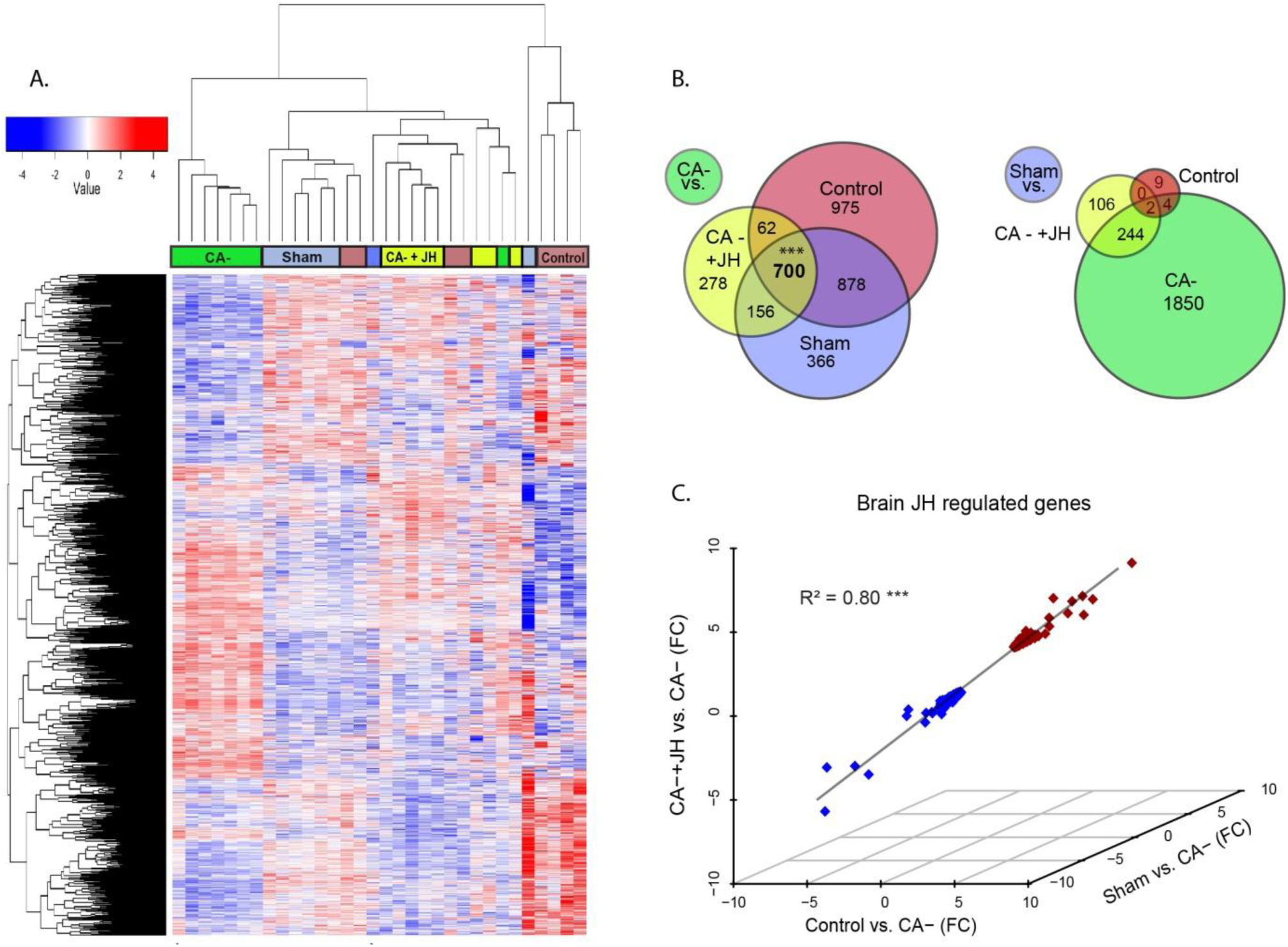
The effect of JH on brain gene expression. Details as in Fig 1 **(A)** Heat map of 3060 DEGs (FDR p < 0.05) in the brain. **(B)** Venn diagrams comparing DEGs between the CA- treatment (left) or the Sham treatment (right) and the three other treatments. **(C)** Three-dimensional correlation analysis for the fold change expression of 700 overlapping DEGs between the CA- and the three other groups. The R^2^ was obtained from a Person correlation test; *** - p < 0.0001 (See Supp. Table 1B for more details).

Only 20 and 15 differentially expressed genes (DEGs) in the fat body and brain, respectively, were found to differ between the control and the sham treatments (Fig 1B right and 2B right) indicating that after five days of recovery the sham operation by itself had only little effect on the patterns of gene expression in these tissues. These findings indicate that the JH replacement treatment, at least partially, recovered the effects of CA removal on the pattern of gene expression in the fat body and this effect cannot be explained by surgery effects.

We next performed Weighted Gene Correlation Co-expression Network Analysis (WGCNA) analyses to identify groups of genes showing a similar co-expression pattern (“modules”) irrespective of whether each gene showed a statistically significant (after FDR correction) difference in expression level. Using this approach, we identified brain and fat body modules that are up- or down-regulated by JH and were differently expressed between bees of the CA- bees and the three other treatments (ANOVA p < 0.05). In the fat body, 10 distinct modules were differently expressed. Modules M3 (886 genes), M8 (249 genes), M9 (244 genes) and M26 (71 genes) were downregulated by JH; modules M2 (1310 genes), M6 (268 genes), M7 (267 genes), M14 (145 genes), M15 (129 genes) and M22 (85 genes) were upregulated by JH (Supp Fig. 1, Supp Table 4a). Four brain modules were also affected by JH: Modules M2 (347genes) and M6 (264 genes) were downregulated by JH, and M9 (196 genes) and M18 (70 genes) were upregulated (Supp Fig 2, Supp Table 4b).

#### Tissue specificity of the JH effect on gene expression

We identified only 156 genes that were affected by JH treatment in both tissues (out of the 1512 and 700 genes differentially expressed between the CA- and all other three treatments in the fat body and brain, respectively). Although this number is 1.37 fold higher than expected by chance (Hypergeometric test p = 1.07E-5, Supp. Table 5) it represents only 11% and 23% of the JH-regulated genes in the fat body and brain, respectively. Most of these genes (139 genes or 89%) were regulated in the same direction producing a significant correlation between the two tissues (Pearson’s correlation: R^2^ = 0.73, t_(df=154)_ = 20.6, p << 0.001). Gene ontology (GO) analyses revealed that the genes upregulated by JH in both tissues are enriched for the GO term “mitochondrion” (fold enrichment (FE) = 3.7, p = 0.033, after FDR correction), and “Citrate cycle” (FE = 13.6, p = 0.058) pathways. No enrichment of any pathway was found for DEGs that are downregulated by JH in both tissues. Seventeen genes (11%) were affected by JH in the opposite direction (e.g., up in the fat body and down in the brain). This analysis shows that although a few processes are similarly regulated in the two tissues, the effect of JH is mostly tissue specific.

#### Pathways regulated by JH in the fat body and the brain

Using KEGG-pathways and GO enrichment analyses (based on DEGs between CA- and CA-+JH treatments that represent the direct effect of JH) we found that JH regulates some of the most important metabolic processes (Table 1; Supp Tables 6a (fat body) and 6b (brain)). Glycolysis, the main pathway for glucose metabolism, was upregulated in both tissues (Fat body: 12/29 genes, FE: 2.5, p = 0.048, Fig 3A right; Brain: 7/29 gene, FE: 4.9, p = 0.046, Fig. 3A left). In the fat body, JH further upregulated the expression of the citrate acid cycle (18/24 genes, FE = 4.5, p = 9.8E-8, Fig. 3A right), Oxidative phosphorylation (OXPHOS, 39/55 genes, FE = 4.26, p = 9.9E-17, Fig. 3B right), biosynthesis of amino acids, and fatty acid biosynthetic process (11/24 genes, FE =3.4, p = 0.02). The citrate acid cycle pathway was not significantly upregulated in the brain when comparing CA- vs CA- +JH, but the difference was statistically significant when we compared the Sham and CA- treated bees (10/24 genes, FE = 3.44, p = 0.03, Supp. Table 6b) suggesting a possible regulation by JH also in the brain. The OXPHOS pathway was not rescued by JH replacement therapy in the brain (7/55 genes upregulated in CA-+JH relative to the CA- bees, Fig. 3B left), but was significantly upregulated in the Sham vs the CA- treatments (24/55 genes, FE = 3.6, p = 1.2E-6). Several catabolic pathways were regulated only in the brain, including downregulation of fatty acid metabolic process (17/60 genes; FE:3.4; p = 0.003).

**Table 1.** KEGG / GO enrichment analyses for JH regulated genes in the fat body and brain. Summary of major pathways and GO terms enriched for genes differentially expressed between the CA- +JH and CA- treatments. Orange background - terms upregulated by JH; light blue - terms downregulated by JH. #of Genes: The number of DEGs in the pathway/term; Fold enrichment: (#DEGs in the pathway / total DEGs) / (#genes in the pathway/background); p-value: Fisher exact test, FDR: false discovery rate correction. The p-values are color-coded from high to low (red to white), WGCNA module/s – the number of the WGCNA module/s which were enriched for the KEGG pathway / GO term and differently expressed between the CA- treatment group and the three other treatments.

| Fat body |  |  |  |  |  |  |
| --- | --- | --- | --- | --- | --- | --- |
|  | KEGG pathway / GO term | # of Genes | Fold Enrichment | P-value | FDR | WGCNA module/s |
| Up | Mitochondrion | 139 | 2.5 | 1.5E-29 | 7.4E-27 | 2, 15, 22 |
|  | Oxidative phosphorylation | 39 | 4.3 | 9.8E-19 | 9.9E-17 | 2, 15 |
|  | Metabolic pathways | 133 | 1.6 | 3.1E-11 | 1.5E-09 | 14, 15 |
|  | Carbon metabolism | 34 | 3.0 | 3.5E-10 | 1.2E-08 | 14, 15 |
|  | Citrate cycle | 18 | 4.5 | 3.9E-09 | 9.8E-08 | 15 |
|  | Proteasome | 20 | 3.8 | 3.6E-08 | 7.4E-07 | 6 |
|  | Lipid particle | 42 | 2.3 | 1.8E-07 | 5.1E-06 | 14, 15 |
|  | Endoplasmic reticulum | 58 | 1.9 | 9.6E-07 | 2.0E-05 | 2, 6 |
|  | Mitochondrial ribosome | 21 | 3.2 | 1.3E-06 | 2.6E-05 | 2 |
|  | Biosynthesis of antibiotics | 41 | 2.0 | 6.7E-06 | 1.1E-04 | 14, 15 |
|  | Ubiquitin-dependent protein catabolic process | 47 | 2.0 | 4.8E-06 | 3.0E-04 | 2, 6 |
|  | Fatty acid biosynthetic process | 11 | 3.4 | 6.0E-04 | 2.3E-02 | 7 |
|  | Glycolysis | 12 | 2.5 | 4.4E-03 | 4.8E-02 | 14 |
|  | Biosynthesis of amino acids | 15 | 2.2 | 4.2E-03 | 5.2E-02 | 14 |
| Down | Cell junction | 34 | 2.2 | 1.2E-05 | 3.0E-03 | 3, 8 |
|  | Cell cortex | 28 | 2.3 | 4.4E-05 | 7.2E-03 | 8 |
|  | Cytoplasmic region | 29 | 2.1 | 1.2E-04 | 1.1E-02 | 3 |

| Brain |  |  |  |  |  |  |
| --- | --- | --- | --- | --- | --- | --- |
|  | KEGG pathway / GO term | # of Genes | Fold Enrichment | P-value | FDR | WGCNA module/s |
| Up | Lysosome | 10 | 4.5 | 2.0E-04 | 6.8E-03 | 9 |
|  | Glycolysis | 7 | 4.9 | 2.1E-03 | 4.6E-02 | - |
|  | Plasma membrane proton-transporting V-type ATPase complex | 6 | 9.9 | 1.9E-04 | 5.5E-02 | - |
|  | Biosynthesis of antibiotics | 14 | 2.3 | 5.3E-03 | 7.0E-02 | 9 |
|  | Carbon metabolism | 10 | 2.9 | 4.5E-03 | 7.4E-02 | - |
| Down | Cytosolic ribosome | 71 | 9.3 | 3.3E-62 | 1.4E-59 | 2 |
|  | Centrosome cycle | 27 | 4.3 | 6.8E-11 | 1.9E-08 | 2 |
|  | Fatty acid metabolic process | 17 | 3.4 | 2.1E-05 | 3.0E-03 | 2, 6 |
|  | Translational elongation | 8 | 6.3 | 9.8E-05 | 1.2E-02 | 2 |

**Figure 3:**
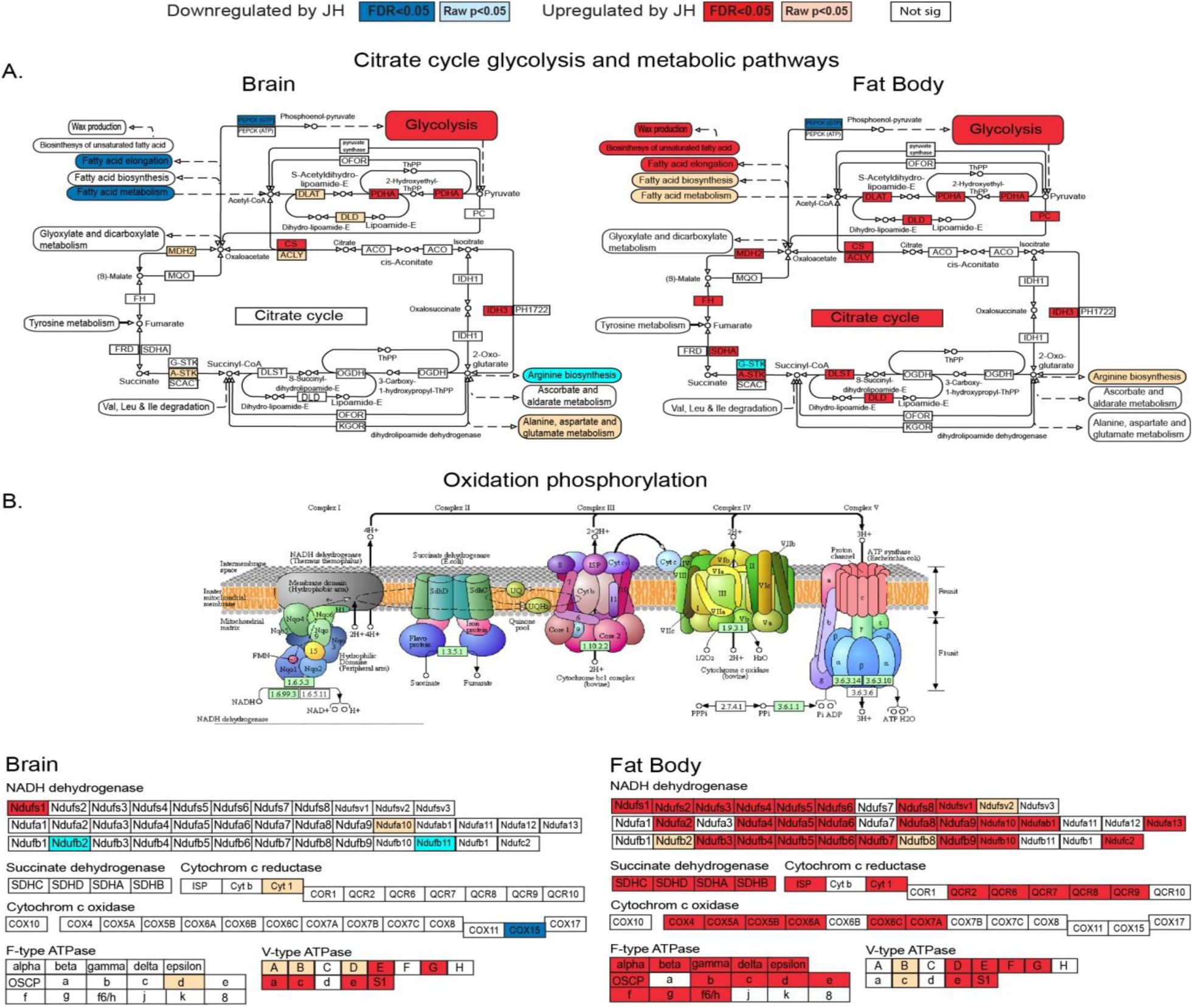
The influence of JH on the expression of genes encoding proteins involved in metabolic pathways. Each gene is depicted as a box with its common name and relative location in the denoted pathways. **(A)** Citrate cycle, glycolysis and related metabolic pathways in the fat body (right) and brain (left). **(B)** The oxidation phosphorylation pathway. The top panel shows a schematic representation of the pathways. The bottom schemes summarise differential gene expression in the fat body (right panel) and the brain (left panel). The effect of JH on gene expression is color-coded: dark blue - downregulation: (FDR p < 0.05); light blue (raw p < 0.05, but p > 0.05 after FDR correction). Dark red - upregulation (FDR p < 0.05); light red (raw p < 0.05, but p > 0.05 after FDR correction). The illustrations are based on the KEGG pathway; citrate cycle: map00020 and OXPHOS: map00190. Full enrichment analysis can be found in Table 1 and at Supp Tables 6a and 6b

The pathway analyses revealed significant effects on the expression of genes involved in protein biosynthesis and degradation (Table 1). The expression of ribosomal protein genes was enriched in both tissues, but the effect was different. In the fat body, we identified an increase in transcript abundance for mitochondria ribosomal (mitoribosome) proteins (21/49 genes; FE = 3.2; p = 2.6E-5; Fig. 4A, right panel). This finding is consistent with the upregulation in OXPHOS and citrate acid cycle because their proteins are synthesized in the mitochondrion by the mitoribosome. The endoplasmic reticulum, the complex responsible for protein folding and packing was also upregulated by JH in the fat body (58/232 genes, FE = 1.9, p = 2.0E-5). The pattern was different in the brain: The expression of cytosolic ribosomal proteins, which were not enriched in the fat body (FE = 0.31; p = 0.41, Fig. 4A, right panel), showed a remarkable enrichment in the brain with an almost universal, downregulation (71/84 genes; FE = 9.3; p = 1.4E-59, Fig. 4A, left panel). Genes encoding translational elongation proteins were also downregulated in the brain (8/15 genes; FE = 6.3; p = 0.01). Taken together, these findings suggest that high JH levels significantly reduce protein production in the brain. On the other hand, there was no significant enrichment for the mitoribosome in the brain (FE = 0.4; p = 0.23, Fig 4A left panel). JH also regulated the expression of genes of the proteasome, a protein-degrading complex that plays a major role in determining protein abundance in the cell. Similar to the ribosome, the expression of proteasomal proteins were regulated in an opposite direction in the two tissues: There was a consistent upregulation in the fat body (20/32 genes; FE = 3.7; p = 7.4E-7; Fig. 4B, right panel) and a statistically not significant enrichment for genes downregulated (after FDR correction) in the brain (6/32 genes, FE = 2.4; FDR p = 0.28, Fig. 4B, left panel). However, it is notable that ten additional brain proteasomal genes were downregulated by JH, but these differences are not statistically significant after FDR correction (Fig. 4B, light blue color in the left panel). Notably, none of the proteasome genes showed upregulated expression in this tissue. Finally, genes encoding proteins of the lysosome, the organelle responsible for biomolecule (including protein) degradation and organelle recycling, were significantly upregulated in the brain (10/45 genes, FE = 4.5, p = 0.007).

**Figure 4:**
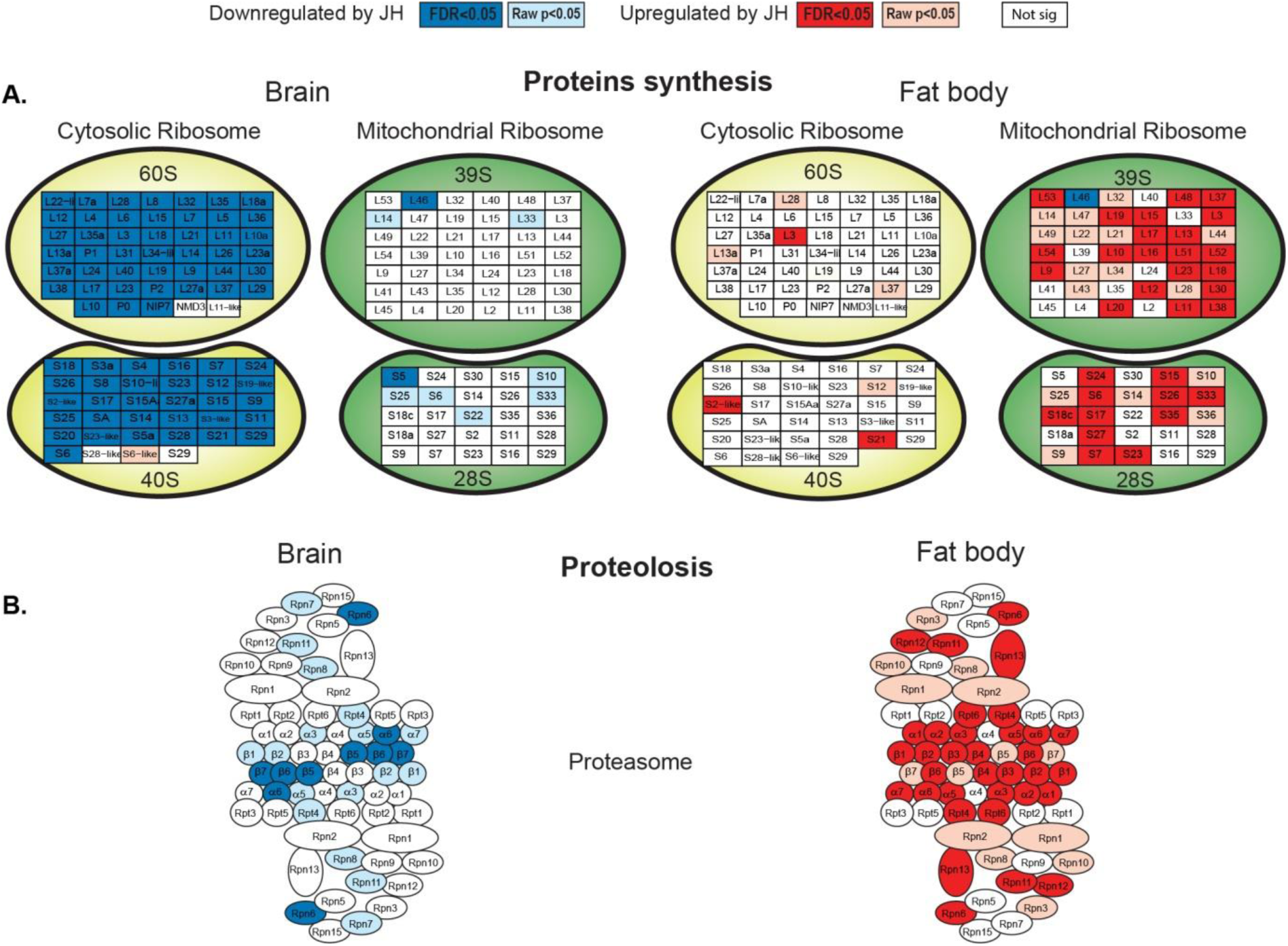
The influence of JH on gene pathways involved in protein synthesis and breakdown. **(A)** Ribosomal protein genes in the cytosol and mitochondria in the fat body (right) and brain (left). Each ribosomal protein gene is depicted as a box with its common name and is assigned to the corresponding ribosomal unit (depicted as ellipsoid structures). **(B)** Genes encoding proteasomal proteins. Each proteasomal protein is depicted as an ellipsoid with its common name. The effect of JH on gene expression is color-coded as in Fig. 3. The illustrations are based on the KEGG pathway; ribosome: map 03010 and proteasome: map 03050 (The proteasome is built of two identical subunits and each protein is drawn twice in the figure).

The WGCNA analysis, which links more genes to JH regulation, extends and supports these findings. Fat body modules 2, 6, 14 and 15, which are upregulated by JH, were enriched for the metabolic pathways: citrate cycle, oxidative phosphorylation (M2 and M15) and Glycolysis (M14), and for the KEGG pathways proteasome and mitochondrial ribosome (M2 and M6, Table 1; Supp. Table 4a). In the brain module 2 was downregulated by JH and highly enriched for the cytosolic ribosome pathway (Supp. Table 4b). Taken together, these analyses suggest that high JH decreases protein biosynthesis in the brain and increase metabolic pathways in the fat body.

#### The effect of JH on the expression of JH pathway genes and vitellogenins

*Krüppel homolog-1* (*Kr-h1*), an established JH readout transcription factor, was on the top of the list of JH upregulated genes showing a seven-, and a four-fold increase expression in the fat body and brain, respectively (CA-+JH vs CA-). Several additional JH signaling genes were also regulated by our JH manipulations (CA-+JH vs CA-; p < 0.05 after FDR correction). These include the putative JH receptor Methoprene-tolerant (*Met*, LOC100647695) which was downregulated by JH in both tissues, and several insulin-pathway genes (e.g., *insulin-like growth factor 1* (LOC100648980); the insulin receptor *chico* (LOC100644779); the insulin binding protein *ltl* (LOC100649210); see Supp Tables 2 and 3). We identified orthologs for all four Vitellogenin (*Vg*) paralogs previously described for the honey bee (Salmela et al., 2016, Supp. Fig. 3A). Vitellogenin (*Vg*) is a conserved yolk protein that is upregulated by JH in most insects but shows a complex and variable interaction with JH in social insects such as honey bees and ants. The predicted *Vg* transcript was the second most abundant transcript in the fat body (4.9% of all transcripts in the control group) and is upregulated by JH (22.5 fold in CA-+JH vs CA-) with over 12K, 24K and 35K counts per million (cpm) in the CA-+JH, Control, and the Sham groups, respectively, but only 600 cpm in the CA- group. *Vg* transcript abundance was much lower in the brain, but it seems to be similarly upregulated by JH (Supp. Fig. 3B left). Fat body *Vg* transcript abundance was much higher compared to the other three *Vg*-like paralogs. Its overall high expression in the fat body and upregulation by JH support the premise that this *Vg* paralog is the major yolk protein precursor in bumble bees. The *Vg like A* paralog was not detected in the brain and was downregulated by JH in the fat body (Supp. Fig. 3B middle left). The *Vg like B* paralog was not effected by JH in any of the tissues (Supp. Fig. 3B middle right). The *Vg like-c* paralog expression was opposite to *Vg:* Its transcript is more abundant in the brain compared to the fat body and is downregulated by JH (Supp. Fig. 3B right). Our findings here for *Vg* and *Kr-h1* were similar to previous studies (Shpigler et al., 2014; Shpigler et al., 2010) providing validation for our RNAseq analyses.

### Transcripts differentially expressed in the brain of dominant and subordinate queenless workers

The results reported above for hormone manipulations are consistent with results we obtained for brain transcriptomes of behaviorally dominant (α) and subordinate (γ) queenless workers, which naturally have high and low JH titers, respectively (Bloch, Borst, et al. 2000). Although none of the DEG differs significantly after FDR correction (Supp Table 7), the 613 transcripts that were differentially expressed (p < 0.05, before FDR correction) show interesting similarities to the effect of JH in the manipulation experiment. First, more transcripts show a trend of lower abundance (361, 59%) in the dominant (high JH) individuals. Second, genes downregulated in the dominant bees were enriched for three pathways related to protein processing, “Protein export” (8/16 genes FE = 9.8, FDR p = 0.0003) “Protein processing in endoplasmic reticulum” (14/88 genes FE = 3.1, FDR p = 0.005) and “cytosolic ribosome proteins” (14/84 genes, FE = 4.34, p = 0.003, Fig. 5A, Supp Table 8). Third, all of the fourteen cytosolic ribosomal proteins that showed a trend towards downregulation in the dominant individuals (Fig 5A), were significantly downregulated also by JH treatment (CA- vs. three control groups). Although the interpretation of this experiment is difficult because of the lack of differences significance after FDR correction, the observed trends are consistent with the premise that some processed that we identified as down-regulated by JH in the manipulation experiment, are similarly downregulated in the brain of workers with naturally high JH titers.

**Figure 5:**
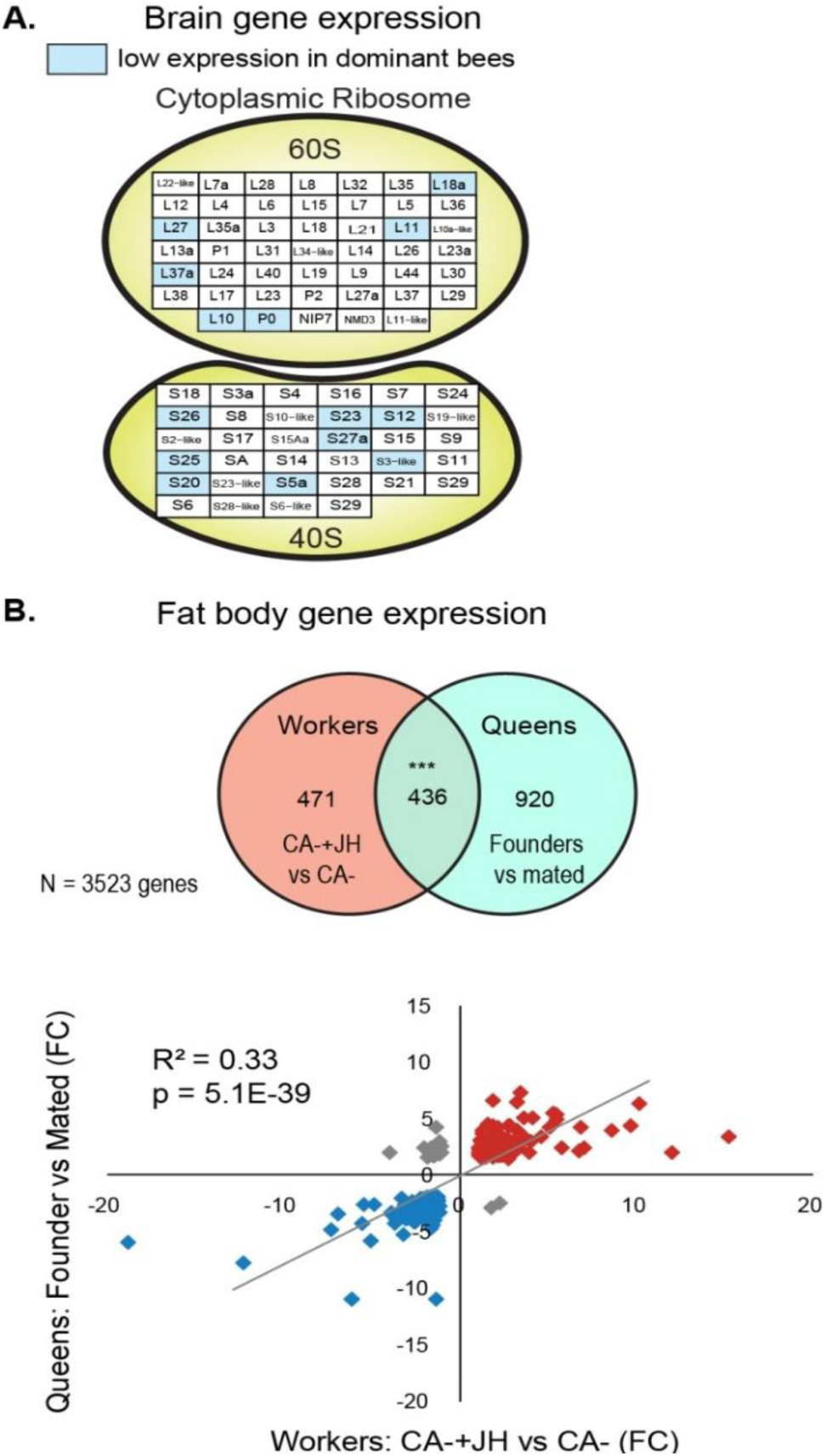
Differential gene expression in bumble bees naturally differing in JH titers. **(A)** Scheme of the cytosolic ribosome (details as in Fig. 4) in the brain of dominant compared to subordinate orphan workers which are assumed to have high and low JH levels, respectively. Transcripts of genes that were differentially expressed before FDR correction (row p-value < 0.05) are marked in a light color (light blue – lower in dominant individuals). None of the genes show statistically significant differences after FDR correction. **(B)** Comparison of fat body transcripts regulated by JH in workers (CA-+JH vs CA-) and fat body transcripts differential expressed between founders and mated queens that are assumed to have high and low JH titers, respectively (queen data from: Amsalem, et al. 2015). Upper panel: Venn diagram of differentially expressed genes between queen’s life stage and workers with manipulated JH levels. Lower panel: Correlation of expression level for 436 fat body transcripts differentially expressed in the two data sets. Each diamond represents one DEG, upregulation is depicted in red, downregulation in blue, opposite direction in grey. The R^2^ was obtained from a Pearson correlation test; *** - p < 0.0001.

### JH regulated genes in the worker fat body are also differentially expressed between reproductive and non-reproductive queens that naturally differ in JH titers

To further explore the links between our hormone manipulation and JH-mediated reproductive physiology, we analyzed data from a database of *Bombus terrestris* queens in which JH is involved in reproduction physiology (Röseler and Röseler 1986, 1988). We compared our list of fat body JH-regulated transcripts with a dataset of DEGs in the fat body of queens at various lifecycle stages (young mated, and egg-laying queens; (Amsalem, et al. 2015)). We identified in our dataset 3523 out of the 4127 genes reported in the queen’s study, 901 of which were regulated by JH in our data (CA- + JH vs CA-), and 1356 between young and egg-laying queens. The overlap of 436 genes between these two gene sets was statistically significant (hypergeometric test p < 0.001; Fig. 5B top). The expression of these common genes was positively correlated (Person correlation test: R^2^ = 0.33, p = 5.1E-39, Fig. 5B bottom). The gene expression profile of reproductive queens is overall similar to that of workers with high JH levels (CA-+JH), whereas that of young mated queens is similar to workers with low JH levels (CA-). This set of overlapping DEGs were enriched for several KEGG pathways including: “mitochondrion”, “carbon metabolism”, “proteasome”, “oxidation phosphorylation”, and “citrate cycle”. These findings are consistent with the premise that JH regulates similar genes and molecular processes in the fat body of queens and workers (Supp Table 9).

### JH regulates different genes in the brains of bumble bee and honey bee workers

Given that in honey bees high JH titers are not associated with ovary activation and reproduction, but rather with the foraging activity of sterile workers, we hypothesized that JH regulates different processes in the brain of the two species. To address this hypothesis, we compared our findings to a microarray study in which honey bees were treated with the JH analog methoprene (Whitfield et al., 2006). We identified in our dataset 2595 of the 3065 probes spotted on the honey bee microarray (Supp Table 10). Out of this set, 454 transcripts in the brains of honey bees and 427 in bumble bees are regulated by JH. The effect of JH treatment differed between the two species. Whereas in the honey bee 62% of the DEGs were upregulated by the JH analog, in the bumble bee, 66% of the DEGs were downregulated (Z-test, Z = 7.7, p < 0.001). Ninety-nine genes were regulated by JH in both species which represent a significant overlap (Hypergeometric test, p = 0.0006, Fig. 6A). Sixty-eight of these genes were regulated in the same direction and 31 in the opposite direction (i.e., up in one species and down in the other, χ^2^ test for independence, p = 5.9E-5). The genes in agreement were enriched for the KEGG pathway “oxidative phosphorylation” (4/27 genes, FE = 10.1; FDR p = 0.06, including genes of the v-type ATPase) suggesting that JH has the same effect on this pathway in both species. The commonly downregulated genes were not enriched for any pathway. The oppositely regulated genes were not significantly enriched to any GO term, but include the *Vg like-C* which was downregulated by JH in the bumble bee (Supp Fig. 3B) and upregulated by methoprene treatment in the honey bee.

**Figure 6:**
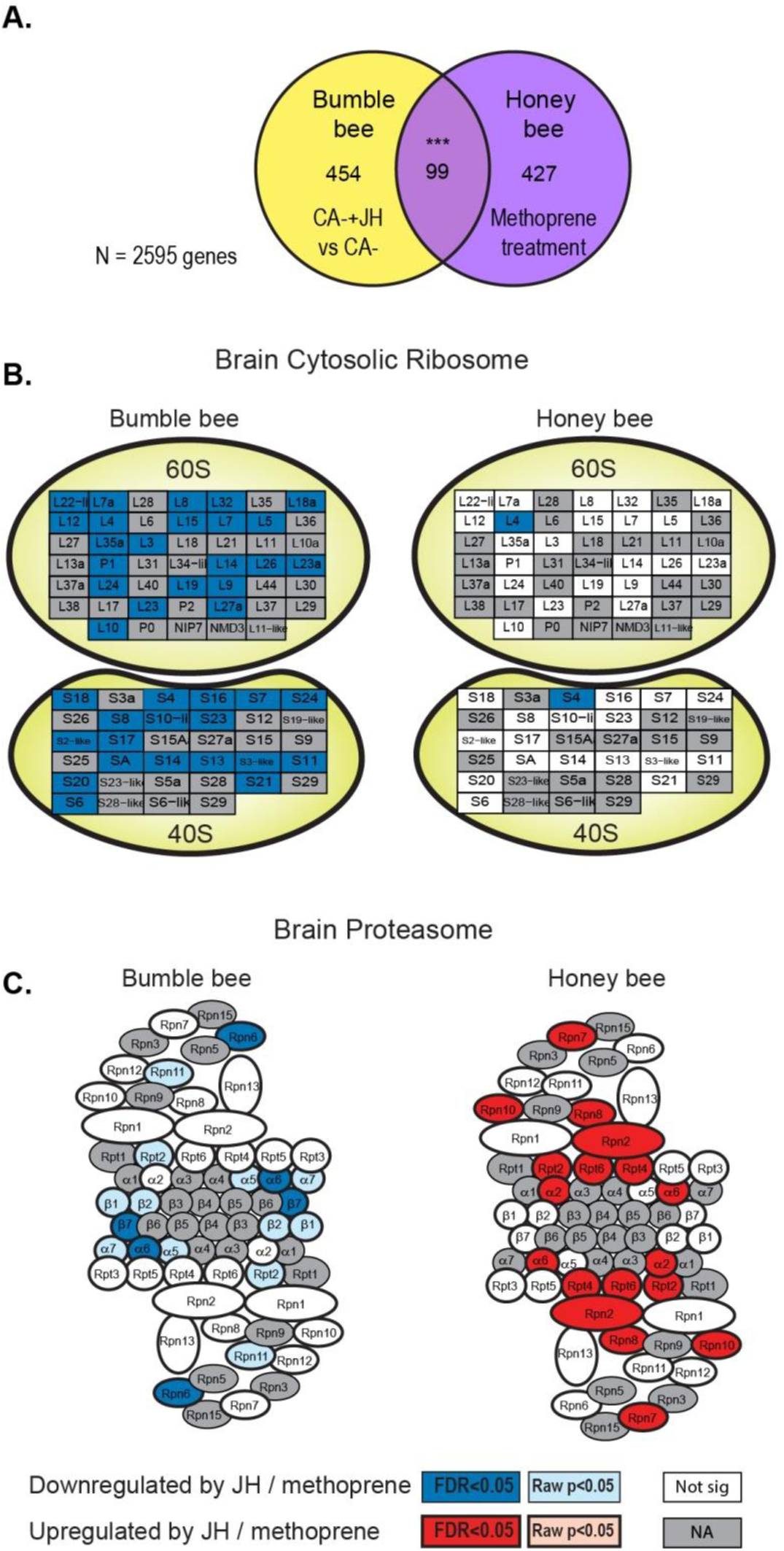
JH regulated genes in the brains of bumble bee compared with honey bee workers. **(A)** A Venn diagram comparing the sets of JH regulated genes in the bumble bee and JH analog (methoprene) regulated genes in the honey bee brain (data from: Whitfield, et al. 2006). **(B)** A scheme summarizing JH influence on the cytosolic ribosomal protein genes. **(C)** A scheme summarizing JH influence on the proteasomal protein gene. Details as in Fig. 4 but the analyses were limited to genes that were found in both data sets. Other details as in Fig. 4; grey staining – a ribosomal protein encoded by a gene not found in the common data set.

JH regulated different genes and pathways in the two species. Most of the JH regulated pathways that we identified above for the bumble bees, using a larger set of DEGs, were also enriched here with a smaller set of DEGs. These include upregulation of the KEGG pathways “Lysosome” (6/20 genes; FE=3.7, p = 0.004) and “Glycolysis” (4/14 genes; FE=9.7, p = 0.08), and downregulation of the “Cytosolic ribosome” (28/34 genes; FE=8.6, p = 7.1E-21, Fig. 6B). None of these pathways were enriched in the set of genes regulated by JH analog treatment in honey bees. Rather, the significantly enriched KEGG pathways include “Oxidative phosphorylation” (11/27 genes; FE=3.9, p = 0.008) and “Proteasome” (8/23 genes; FE=3.4, p=0.15), which were upregulated, opposite of the pattern in bumble bees (Fig. 6C). No pathways were significantly downregulated by JH analog in the honey bee. These analyses suggest that by contrast to the bumble bee, the expression of pathways involved in protein production and turnover is not downregulated by JH in the honey bee brain.

## Discussion

Our transcriptomic analyses show that in bumble bee workers JH has diverse and mostly tissue-specific effects in the brain and the fat body. JH activates the mitochondrion and additional metabolic and biosynthetic processes in the fat body, an organ pivotal in energy storage and utilization in insects (Arrese and Soulages 2010). These findings are in line with earlier studies showing that JH activates the ovaries and several exocrine glands (Shpigler, et al. 2014) indicating that JH is the major gonadotropin of the bumble bee. The effects are different in the brain in which high JH downregulates the expression of many genes showing significant enrichment for pathways regulating protein turnover such as ribosomal, and proteasomal proteins. JH also upregulates the lysosome that recycles proteins and organelles. Similar downregulation of pathways controlling protein turn-over and increase in lysosome activity is commonly associated with aging or stress in diverse organisms (Ryazanov and Nefsky 2002; Marion, et al. 2004; Lim and Zoncu 2016), but have never been shown to be down-regulated by increase fertility. Decreased protein biosynthesis may compromise brain maintenance and processes such as learning and long-term memory (Hernandez, et al. 2015). These JH effects in the bumble bee brain point to a previously unrecognized cost of gonadotropic hormones to the brain in the form of decrease protein biosynthesis and increase organelle recycling. We further suggest that this cost would have been maladaptive to highly eusocial honey bees which maintain exceptionally high fertility over extended periods lasting over several years. Consistent with this premise, we did not find evidence for a similar effect of JH on brain gene expression in the honey bee in which JH does not function as a gonadotropin in adult females

We found that most *B. terrestris* fat body DEGs are upregulated by JH. The upregulated genes are enriched for mitochondria genes that include key metabolic pathways such as citrate acid cycle, glycolysis, and oxidative phosphorylation, suggesting that as in other insects, JH activates the bumble bee fat body (Glinka and Wyatt 1996; Panaitof and Scott 2006; Yamamoto, et al. 2013). Upregulated genes were also enriched for mitoribosomal proteins which are necessary for translating the JH regulated transcripts into proteins. The fat body transcriptome analysis is consistent with JH upregulating ATP production which is necessary to support the high energetic demands of the activated reproductive system, and the biosynthesis of yolk proteins (Arrese and Soulages 2010). Indeed, we found that the fat body expresses large amounts of the *Vg* transcript, accounting for about 5% of the total transcripts in the fat body of bees with high JH titers. To test if the fat body is similarly activated in bees in which JH increases naturally, we compared our findings to a transcriptomic database of bumble bee queens which naturally vary in JH titers. We found that the set of fat body transcripts that we found to be regulated by artificial manipulation of JH titers in the worker fat body (Fig. 2) significantly overlap with genes differentially expressed between egg-laying founders bumble bee queens and pre-diapause mated gynes which naturally have high and low JH titers, respectively (Amsalem, et al. 2015). Thus, as in other insects (Stepien, et al. 1988), JH-regulated reproduction in the bumble bee is associated with a significant increase in energy demand and resource utilization. Elevated metabolism in reproductive females is not unique to bumble bees. For example, in mammals mitochondria activity is elevated in the trophoblast and placenta of pregnant females (Van Blerkom 2009; Ramalho-Santos and Amaral 2013). This evidence for the high metabolic cost of JH raises the question of how can the body meet these increasing energy demands of multiple tissues? One common way for contending with this challenge is to store sufficient energy before the reproductive season. Our findings suggest an additional, not mutually exclusive mechanism, in which some tissues pay a cost by reducing metabolism or biosynthetic activity.

By contrast to the fat body, in the brain, overall more genes were downregulated by JH. The pathways showing the most consistent down-regulation are the cytosolic ribosome and translation elongation. This transcriptomic signature suggests that high JH levels downregulate protein biosynthesis in the brain. Fatty acid metabolism pathways were also downregulated. The brain uses fatty acids as metabolites and also as building blocks of cell and organelles membranes (Tracey, et al. 2018); deficiency in fatty acid impairs learning abilities in honey bees (Arien, et al. 2015). On the other hand, genes encoding proteins of the lysosome and phagosome were upregulated by JH. The lysosome is central for degradation and recycling of organelles and macromolecules, including proteins (Appelqvist, et al. 2013; Settembre, et al. 2013). The increased expression of genes encoding lysosomal and phagosomal proteins thus suggests that JH activates the degradation of available organelles, which can be then reused as building blocks for essential proteins and lipids. Similar upregulation of lysosomal protein genes and downregulation of ribosomal protein genes is associated with conditions of starvation, stress, and aging in other species (Marion, et al. 2004; Settembre, et al. 2013; Mony, et al. 2016). Do similar transcriptomic changes exist in the brain of bees in which JH titers increase naturally? We addressed this question by studying orphan workers in which JH levels are positively correlated with dominance rank (Bloch, et al. 1996; Bloch, Simon, et al. 2000). Although the interpretation of this experiment is somewhat difficult because none of the 613 differentially expressed transcripts were statistically significant after FDR correction for multiple comparisons, there was a very clear and consistent trend. The cytosolic ribosome proteins expression in the brain of the dominant bees, that typically have high JH titers, were downregulated showing a response that is overall consistent with the pattern we found in workers for which we manipulated JH titers.

Our findings are consistent with additional evidence for a cost for a gonadotropic JH in insects (for a recent review see (Rodrigues and Flatt 2016). For example, *Drosophila* flies treated with a JH analog show reduced lifespan (Yamamoto, et al. 2013) and compromised immune functions (Schwenke, et al. 2016; Schwenke and Lazzaro 2017). JH also appears to mediate a trade-off between fecundity and life span in *Polistes* paper wasps (Tibbetts and Huang 2010; Tibbetts and Crocker 2014). In butterflies individuals treated with a JH analog showed reduced learning performance (Snell-Rood, et al. 2011). The evidence of costs of high JH titers in insects is reminiscent of the costs of testosterone (and other androgens), the male gonadotropic hormone of vertebrates (Wingfield, et al. 1990; Wingfield 2017). These apparent costs of high gonadotropic hormone levels raise the question of how can the exceptionally fertile queens of highly eusocial species such as honey bees keep on reproducing over extended periods without paying these costs? Or in other words, how do highly eusocial insects defy the evolutionary ancient trade-off between reproduction and maintenance/ longevity? We hypothesized that maintaining such exceptional fertility over long periods required evolutionary modifications in hormonal signaling pathways which alleviate the ancient cost of high JH titers (Bloch, et al. 2009). We tested this hypothesis by re-analyzing a previously published dataset of brain gene expression of honey bees treated with a JH analog. Consistent with the idea that JH signaling was modified, we found that in the honey bee, in which JH is not a gonadotropin, high JH titers were associated with an overall up-, not down-regulation of brain gene expression and with no down-regulated expression of ribosomal proteins or other pathways involved in protein synthesis. These findings are consistent with evidence that JH treatment has a positive effect on some learning tasks in honey bees (McQuillan, et al. 2014). The influences of JH on the fat body also differ substantially between the honey bee and the bumble bee. Whereas in the bumble bee JH overall upregulates metabolic and protein synthesis pathways, in the honey bee JH down-regulates these pathways (Lu, et al. 2017). Thus, whereas in bumble bees JH seems to downregulate processes in the brain and activate the fat body, in the honey bee it seems to overall activate the brain while downregulating the fat body.

We speculate that in the ancestral solitary and annual bees, JH functioned as the major gonadotropin activating the ovaries and other tissues related to reproduction while reducing investment in tissues such as the brain (Hartfelder 2000; Smith, et al. 2013; Kapheim and Johnson 2017). We further assume that the cost of high JH is manageable for these short-lived, low fertility, bees. The evolution of sociality was associated with the development of a reproductive skew in which some females (i.e., queens) became significantly more fertile compared to solitary bees (Michener 1974). Nevertheless, species with an annual life cycle apparently could still sustain relatively short-term costs because the life expectancy of reproductive queens is relatively low and they increase their fitness by sacrificing long-term maintenance for optimizing fertility during their limited reproductive period. The evolution of division of labor can further mitigate these costs because the cognitively demanding foraging activities are performed by individuals with low JH titers and undeveloped ovaries as typical to bumble bees (Cameron and Robinson 1990; Shpigler, et al. 2016). These costs, however, are not bearable for long- lived highly fertile queens which are a hallmark of advanced eusociality. The transition to this stage in the evolution of sociality required decoupling the reproduction-maintenance/longevity tradeoff. Although one can speculate on several solutions for reducing this cost (e.g., modifications only in JH signaling in the brain), it appears that in honey bees and some ants the evolution of JH signaling involved substantial reduction in the gonadotropic functions of JH, perhaps because ancestrally JH regulated life history switches between reproductive and non-reproductive stages that affect many tissues in the body (Rodrigues and Flatt 2016). When JH was freed from its role as the major coordinator of reproductive tissues, it can regulate (old or new) functions such as age-related division of labor and other complex behaviors. Thus, our findings link two remarkable physiological traits of advanced eusocial insects: first, they defy the widespread trade-off between reproduction and longevity, second, JH, which is the ancient insect gonadotropin, does not appear to regulate fertility in some social insects such as honey bees and some ants - a riddle posed about three decades ago but still has no satisfactory answer (Cameron and Robinson 1990; West-Eberhard and Turillazzi 1996). Our findings suggest that these two traits are linked because high JH titers would have caused a serious cost to queens that are highly fertile over extended periods, and therefore the evolution of eusociality in these lineages was associated with modifications in JH signaling pathways.

## Acknowledgments

We thank Yafit Brener, Gal Hadad and Adam J Siegel for assistance with bee collection, Nadav Yayon for manufacturing the surgical table. This work was supported by research grants from the US–Israel Binational Agricultural Research and Development (BARD) fund (IS-4418-11 to G.B., G.E.R, and M.B), the Vaadia-BARD Postdoctoral Fellowship (award No. FI-462-2012 to H.Y.S), and the Israeli Ministry of Science Yitzhak Shamir fellowship (to H.Y.S)

## Supplementary material

### Supplementary table and figures

**Supp Table 1.** JH replacement therapy rescued the expression level of hundreds of genes in the brain and fat body. Person correlation test results for shared genes differently expressed between the CA- group and the other three experimental groups: Fat body **(A)** Brain **(B)**. The number are R^2^ and *** represent p ≪ 0.0001.

| **A. Fat body (1512 genes)** | **Sham vs CA-** | **CA-+JH vs CA-** |
| --- | --- | --- |
| **Control vs CA-** | 0.783 *** | 0.522 *** |
| **Sham vs CA-** | - | 0.662 *** |

| **B. Brain (700 genes)** | **Sham vs CA** | **CA-+JH vs CA-** |
| --- | --- | --- |
| **Control vs CA-** | 0.882 *** | 0.808 *** |
| **Sham vs CA-** | - | 0.846 *** |

*Supp Excel files (not included in the initial submission)*

**Supp Table 2: The effect of JH on bumb lebee workers fat body gene expression**

**Supp Table 3: The effect of JH on bumble bee workers brain gene expression**

**Supp Table 4a: Fat body WGCNA and modules GO analysis**.

**Supp Table 4b: Brain WGCNA and modules GO analysis**.

**Supp Table 5: Brain and Fat body gene expression comparison, CA- vs CA-+JH**

**Supp Table 6a: Fat body KEGG and GO enrichment analysis for JH regulated genes**.

**Supp Table 6b: Brain KEGG and GO enrichment analysis for JH regulated genes**.

**Supp Table 7: Dominant vs subordinate bumble bee workers gene expression**.

**Supp Table 8: Dominant vs subordinate GO enrichment analysis**.

**Supp Table 9: Comparison of fat body gene expression between bumble bee workers and queens (Amsalem et al**., **2015)**

**Supp Table 10: Comparison of JH related genes in bumble bees and honey bees (Whitfield et al**., **2006)**.

**Supp. Fig. 1:**
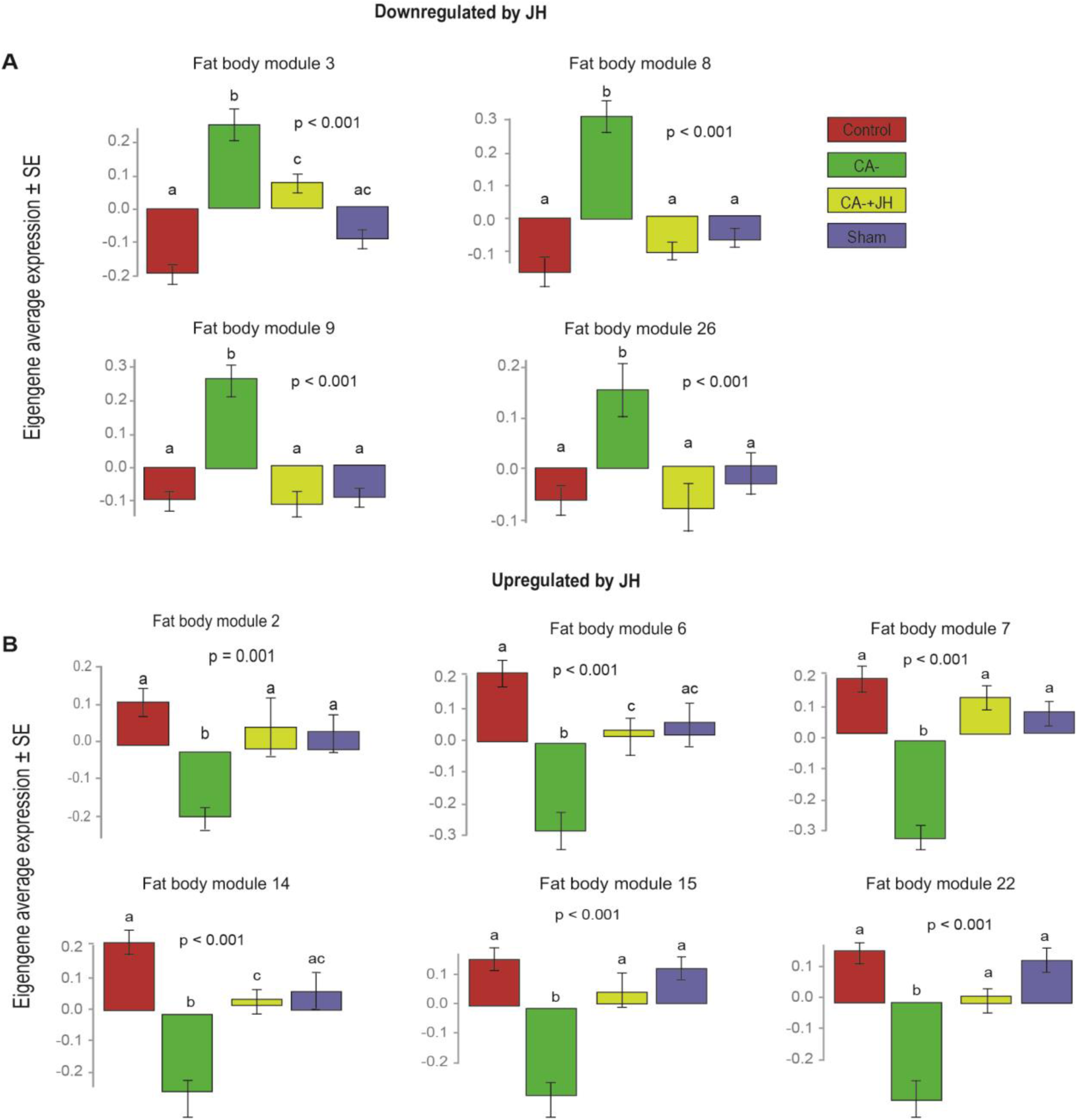
WGCNA analysis of gene expression in the fat body. The bar represents the module eigengene (the best summary of the standardized module expression data) average expression ± SE, for each experimental treatment, control (red), allatectomized (CA-, green), Replacement therapy (CA-+JH, yellow), Sham-operated (blue). **A**. Downregulated by JH: module 3 (886 genes) module 8 (249 genes), module 9 (244 genes), module 26 (71 genes). **B**. Upregulated by JH: module 2 (1310 genes), module 6 (268 genes), module 7 (267 genes), module 14 (145 genes), module 15 (129 genes) module 22 (85 genes). The p-values were obtained from ANOVAs, different letters above bars represent significant differences between the experimental groups in a Limma pairwise comparison. For a summary of all modules see Supp. Table 4a.

**Supp Fig. 2:**
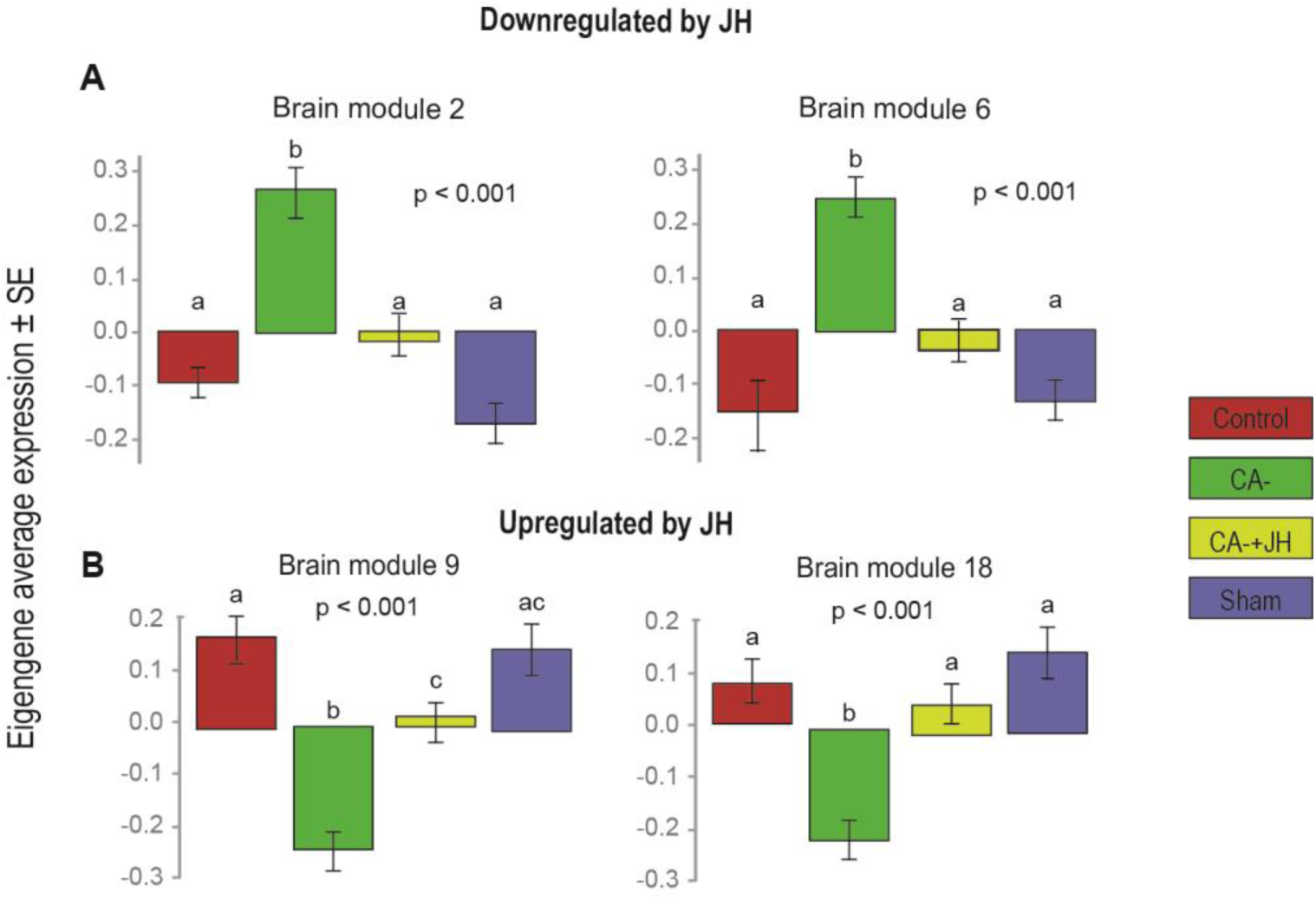
WGCNA analysis of gene expression in the brain. Details as in Supp fig 1. **A**. Downregulated by JH: module 2 (347 genes) module 6 (264 genes). **B**. Upregulated by JH: module 9 (196), module 18 (70). For a summary of all modules see Supp. Table 4b.

**Supp Fig. 3:**
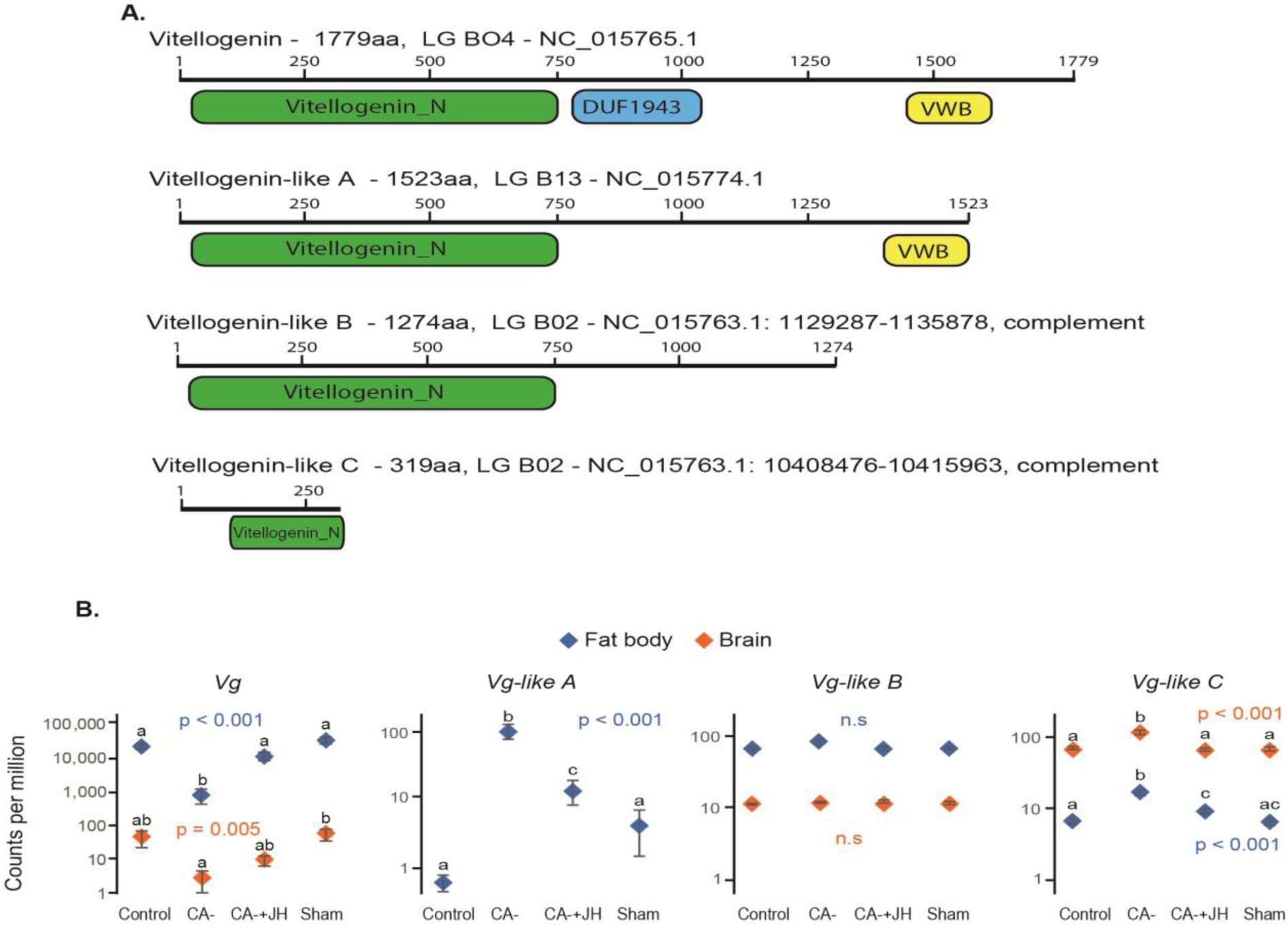
The influence of JH on the expression of Vitellogenins. (A) Gene models of the *Bombus terrestris* four *Vg-like proteins*. (B) Transcript abundance for the four bumble bee *Vg*-like genes. Orange diamonds - brain; blue diamonds - fat body. The *p*-values summarise ANOVA (with FDR correction). Treatments marked with different lowercase letters are significantly different in Limma pairwise analyses (FDR p < 0.05)

## Material and methods

### Bees

For the RNAseq experiments, we collected *Bombus terrestris* workers from eight “donor” colonies. To reduce genetic variation, the donor colonies were founded by full-sister queens (genetic relatedness, r = 0.75). Queen mating and colony initiation were performed by BioBee Biological Systems (Sde Eliyahu, Israel) according to standard rearing techniques. When these colonies contained a queen, 5-10 workers, and brood at all stages of development, they were shipped to the Hebrew University of Jerusalem. The colonies were placed in a wooden nesting box (21 x 21 x 12 cm) with a front wall and cover made of transparent acrylic plastic (Plexiglas™). The nesting boxes with the bees were housed in an environmental chamber (28 ± 1°C; 50 ± 5 % RH) in constant darkness at the Bee Research Facility at the Edmond J. Safra Campus of the Hebrew University of Jerusalem, Givat Ram, Jerusalem. The colonies were kept indoor and were fed *ad libitum* with commercial sugar syrup (70% sugar, purchased from Ployam Pollination Services, Kibbutz Yad Mordechai, Israel) and fresh pollen (collected by honey bees) mixed with sugar syrup. Bee collection and observations were done under dim-red light that the bees do not see well.

### The influence of JH on fat body and brain gene expression

We manipulated JH levels by surgically removing the corpora allata (CA) glands (*allatectomy*), the only source of JH in insects (Riddiford 2008). We used our previous protocol as detailed in (Shpigler, et al. 2014). Briefly, newly emerged worker bees were collected (up to 18 hours after emerging from the pupa, easily recognized by their light body color) from donor colonies. At this age, the cuticle is relatively soft and easy to manipulate. We placed the collected callow bees in a cage (20×20×10 cm) provisioned with *ad libitum* sugar syrup and pollen. For the dissection, we anesthetized the bees on ice for 5-20 min (the variation in chilling duration was due to individual differences in body size, and a consequence of the dissection order as the bees were chilled in groups of four) and when immobile, fixed them under a stereoscopic microscope (Nikon SMZ645, X50) onto an ice-chilled metal stage using modeling clay. The dorsal side of the bee faced up and the head bent down to expose the thin neck cuticle connecting the thorax and the head. We used a fine scalpel to open a latitudinal incision in the posterior part of the head capsule. We then moved the inner membrane and trachea to expose the CA glands. Both corpora allata glands were gently grasped with fine forceps and detached. The entire procedure took between 2-5 minutes, and the cuticle resumed its original shape; the incision appeared to self-seal within a few hours following the operation.

Sham-operated bees (*‘Sham’*) were handled and dissected in a similar way, however, the CA were only touched gently and not detached. Control bees (*‘Control*’) were anesthetized and handled similarly but were not operated. At the end of the operation, the bees were placed in a small wooden cage (12×8×5 cm), with other similarly manipulated bees, and were left to recover overnight in an incubator (32°C, 70% RH). On the second day, the surviving bees from each treatment group were assigned to groups of three, each transferred to a fresh, clean wooden cage with clear glass walls (12×5×8 cm). The average survival rate for the first day in the three experimental groups was 50% for the allatectomized (*‘CA-’*) bees, 80% for the sham, and 100% for the control bees. Bees that survive the first day after operation showed similar survival rates: Survival during days two to five was 86% (45/52) for the CA- bees, 94% (34/36) for the sham-treated bees, and 97% (35/36) for the control bees (Fisher exact test, p = 0.17). Only groups in which all the three bees survived for the whole five days of the experiment were used for the RNAseq analysis. For replacement therapy, (*‘CA-+JH’*) half of the allatectomized groups were randomly chosen and treated as follows. The bees were chilled on ice for 3-5 minutes, and when anesthetized, were treated topically with 70µg of JH-III (Sigma, cat #: J-2000) dissolved in 3.5µl Dimethylformamide (DMF, J.T Backer, cat #: 7032), giving a final concentration of 20 µg/µl. The JH solution was applied to the dorsal part of the thorax. JH treatment was done twice at day 2 and day 4 from emergence. Our previous study using this protocol showed that two tandem JH treatments successfully reverted the effect of CA removal on various JH regulated traits, including ovarian activity, wax secretion and *Vg* expression (Shpigler et al., 2014). On the same two days, we similarly handled and chilled the sham and CA- bees but treated them only with the vehicle (3.5µl DMF). The control bees were chilled on days 2 and 4, but otherwise untreated. Following treatment, the bees were returned to their original cages. The JH treatment did not affect the survival of the bees as the CA-+JH group survival (22/26) was not different from the CA- group (23/26; Fisher exact test, p = 1.0). The bees from all treatment groups were placed in an incubator (28°C ± 0.5°, 70% ± 5% RH) for four days and were monitored daily for survival. On day five from emergence, the bees were collected by flash freezing in liquid nitrogen and immediately transferred to individually marked centrifuge tubes on dry ice. The samples were kept in an ultra-freezer (−80°C) until tissues dissection.

### The influence of dominance rank on brain gene expression

We paint marked the workers of the control groups (see above) and carefully recorded their behavior. We performed two sets of observations (20 min each); the first, at the age of three days, and the second at the age of five days, just before collection. The dominance index was calculated following the method described by (Bloch, et al. 1996). Briefly, for each encounter of a pair of bees, we recorded which bee advanced and which retreated. The dominance index was defined as 1 – retreats / total encounters, and thus, ranges between 0 - 1. The bees were classified according to their dominance rank; the most dominant individual (highest score) in the triplet was dubbed “α,” the median “β,” and the individual with the lowest rank “γ.” For the RNAseq analysis, we used only the α and γ ranked individuals. Eight dominant and seven subordinate five-day-old bees were collected for transcriptomic analysis as described above.

### Tissue dissections and RNA extraction

We chose eight bees from each experimental group (except for the Control group from which we collected 15, see above) for the transcriptomic analysis. We first separated the head of the bee from the rest of the body and then opened small windows in the frontal part of the head capsule. The opened heads were lyophilized for 60 min to facilitate the dissection of the brain. After freeze-drying, we accomplished the removal of the frontal head cuticle, cleaned glandular remains, and removed the whole brain from the head capsule. The brain was cleaned of any other tissue and placed it in a fresh centrifuge tube. All dissections processing was carried out on dry ice to minimize RNA degradation. RNA was extracted using the RNeasy kit Invisorb Spin Tissue RNA Mini Kit (Invitek, Germany) according to the manufacturer’s protocol. RNA samples were shipped on dry ice to the Carver Biotechnology Center at the University of Illinois (UIUC) for RNA sequencing. Bee bodies were sent on dry ice to the lab of Dr. Robinson at the Carl R. Woese Institute for Genomic Biology, UIUC, where the fat body RNA was extracted. The bee abdomen was separated and immersed in chilled RNA-later ICE (Thermo-Fisher, MA, USA), for 16-18 hours. The gut, ovaries and other internal organs were removed leaving the fat body tissue attached to the abdominal cuticle. The fat body was placed in a fresh centrifuge tube. RNA was extracted using the RNeasy kit, Qiagen, (OR, USA), followed by DNase treatment. For both the brain and fat body tissues, 1µg of RNA from each sample was used for whole transcriptome expression analysis. The RNA integrity was determined using a Bioanalyzer 2100 (Agilent).

For sequencing, all libraries were diluted to a 6nM concentration. RNA-Seq libraries were constructed with the TruSeq Stranded mRNA HT (high throughput kit, Illumina cat #: RS-122-2103) using an epMotion 5075 robot (Eppendorf). The libraries were pooled in equimolar concentration as per instructions and each pool was quantitated by qPCR. Paired-end for brain and single-end for fat body sequencing (read length = 100nt) was performed on an Illumina HiSeq 2500 using a TruSeq SBS sequencing kit, v4. FASTQ files were generated with CASAVA 1.8.2. Pooled RNA-seq samples produced an average of 34,013,322 reads per sample for brain tissue and 23,257,243 reads per sample for fat body tissue. RNA-seq files have been deposited in the Sequence Read Archive under BioProject number PRJNA497863.

### Bioinformatics analysis

Sequencing reads were trimmed with Trimmomatic version 0.30 and aligned to the *B. terrestris 1*.*0* reference genome (Elsik, et al. 2014; Sadd, et al. 2015)(Elsik et al., 2014; Sadd et al., 2015) using STAR version 2.4.0 with alignIntronMax option set to 10000. Numbers of reads per gene were counted using featureCounts version 1.4.3 with default settings. EdgeR (Robinson, et al. 2010) was used to normalize using the trimmed mean of M-values (TMM) method. A total of 9,446 and 8,984 genes in the brain and fat body respectively and were tested for differential expression. We used the limma-voom method to calculate a one-way ANOVA followed by pairwise comparison between the groups in R version 3.2.0. P-value correction for multiple testing was done using the false discovery rates (FDR) method (Benjamini and Hochberg 1995). Lists of differentially expressed genes (DEGs) were determined based on FDR < 0.05.

We next performed Weighted Correlation Co-expression Network Analysis (WGCNA) using the R package *WGCNA* (Langfelder and Horvath, 2008) to identify groups (“modules”) of co-expressed genes. The appropriate soft-thresholding power to achieve a scale-free topology was estimated separately for each tissue; fat body plateaued at power = 7 while brain plateaued at power = 12. Genes were assigned to modules using the *blockwiseModules* function with default parameters except for power = 7 or 12, maxBlockSize = 25000, networkType = “signed hybrid” and minModuleSize = 20. Using these parameters, we identified 35 and 47 modules (ordered by the number of genes in each module: M1 > M2) in the fat body and brain respectively. The expression pattern of each module was summarized by calculating an eigengene value for each sample, which were then tested for differential expression in a similar fashion to individual genes: a limma model was used to calculate a one-way ANOVA test followed by pairwise comparisons. FDR correction was applied to each set of 35 or 47 modules

For Gene Ontology (GO) a reciprocal BLAST was used to create a one to one orthologous gene list from *Bombus terrestris* to *Drosophila melanogaster*. This list included 5214 fly genes ids (Supp Table 2 and 3) and these were used as background lists for the GO and KEGG pathway enrichment analysis. We further looked separately on pathways enriched in DEGs that are up- or down-regulated. The GO and KEGG pathway enrichment was done in DAVID 6.8 (Huang, et al. 2009). For the analysis, we used the standard parameters and the GO.FAT option. The enrichment was calculated based on the frequency of significantly differentially expressed genes (after FDR correction) in a pathway compared to the expected frequency based on the background gene list used in the analysis. The significant of the enrichment was calculated using Fisher exact test followed by Benjamini fold discovery rate correction in DAVID.

For KEGG pathway enrichment and figures, pathway information for *Bombus terrestris* was downloaded as a single *keg* file from the KEGG website. A list of genes for each pathway was derived from this keg file. DEGs were matched to each pathway and enrichment analysis was performed in R. Enrichment was calculated based on the frequency of significantly differentially expressed genes in a pathway compared to the expected frequency based on the total gene set used in gene expression analysis. The hypergeometric test was calculated using the hyperfunction from the ‘stats’ package in R.

The KEGG pathway Ribosome (ko#: 3010) includes genes from the cytoplasmic and the mitochondrial ribosomes (mitoribosome) together, for the analysis, we split between the two ribosomes based on the NCBI annotation of the genes. Many of the *B. terrestris* mitoribosome genes are not included in the KEGG dataset. However, in the NCBI dataset, 33 additional mitoribosome genes are annotated, making this pathway more reliable as the mitoribosome includes 80 genes in most eukaryotes (Greber and Ban 2016). For the analysis of this pathway, we used both the KEGG and the NCBI annotated genes and included all the genes in the figures.

For the analysis of the dominance\ subordinate gene expression trend, we used the raw p-value for our analysis as none of the genes were statistically significant after FDR correction. We split the trending genes list to up and down-regulated in dominant bees compare to subordinate bees and checked for GO and KEGG enrichment using DAVID as explained above.

### Vitellogenins analysis

Salmela, et al. (2016) identified four honey bee *Vitellogenins* proteins *(amVg, amVg like-A, amVg like-B, and amVg like-C)*. Using BLAST we identified four *B. terrestris* orthologs, each located on a different linkage group. All four predicted *Vg*-like proteins contain a *Vitellogenin* domain, or at least part of it (Supp Fig 2A). The *B. terrestris* LOC100650436 is predicted as the ortholog of *Vg* protein and most similar to the protein that accumulates in the developing eggs of honey bees and other insects and functions as the egg yolk precursor (Tufail and Takeda 2008; Salmela, et al. 2016). The other three honey bee *Vg* paralogs were also identified in the *B. terrestris* genome: LOC100643258 gene similar to *amVg like-A*, LOC100644917 similar to *amVg like-B*, and LOC100649251 similar to am*Vg* like-C, and were designated btVg-like-A, btVg-like-B, and btVg-like-C, respectively

### Comparison to published data

We compared our data with published data sets from bumble bees and honey bees. Gene expression data sets were obtained from the supplementary tables of the following studies: Transcriptional profiling (RNAseq) of *B. terrestris* queens fat body at various life stages from (Amsalem, et al. 2015); whole-brain transcriptional profiling (microarray) of methoprene treatment of honey bee workers from (Whitfield, et al. 2006). Overlap of statistically significant differentially expressed genes from each study was tabulated and hypergeometric tests were performed to test whether the overlap of the two datasets is statistically different from expected by chance.

